# Precise Instruction and Consideration of the Vertical *and* Horizontal Force Component Increase Validity and Reliability of the 90:20 Isometric Posterior Chain Test

**DOI:** 10.1101/2024.06.06.597848

**Authors:** Dominic M. Rasp, Florian K. Paternoster, Jan Kern, Ansgar Schwirtz

## Abstract

Hamstring injuries are associated with decreased hamstring strength. Matinlauri et al.’s 90:20 Isometric Posterior Chain Test (90:20 IPCT) efficiently assesses hamstring strength, but has not been validated so far. Furthermore, their rather unprecise original instruction allows high variability in test execution. We added a new instruction and parameters and examined, whether this measure leads to increased reliability and validity.

We assessed hamstring strength of 23 sport students via the 90:20 IPCT under the original instruction, to exert vertical force, and our new instruction, to exert vertical *and* horizontal force. Instead of only using bare vertical force as outcome parameter under the original (Fz_V) and our new instruction (Fz_VH), we also calculated the resultant force (Fres_VH) and the applied torque onto the force place (M_F_ortho_VH). To test for validity, we correlated the outcome parameters with peak torque of gold standard dynamometry. Furthermore, we measured muscle activities of the mm. rectus femoris, biceps femoris, semitendinosus, and gluteus maximus under our new instruction and compared them to those under the original parameter (Fz_V) via one sample t-tests. To evaluate reliability, tests were repeated on two separate days, for which we calculated intra class correlation coefficients (ICCs) and coefficients of variation (CVs).

Our new instruction and parameters (Fz_VH, Fres_VH, M_F_ortho_VH) showed better validity (mean r = 0.77, r = 0.81, and r = 0.85) and equally good or better reliability (ICCs: 0.87, 0.89, and 0.94; CVs: 4.7 %, 4.1 %, and 4.7 %) than the original instruction and parameter (Fz_V) (mean r = 0.70; ICC: 0.91; CV: 5.6 %). There were no differences in muscle activities between the parameters and instructions of the 90:20 IPCT.

We recommend our new instruction and the applied torque onto the force plate as it makes the 90:20 IPCT a more reliable and valid tool to assess hamstring strength.

## Introduction

The proportion of hamstring injuries (HSI) in all injuries in soccer and the resulting absent time caused by HSI has doubled over the last 20 years [1]. A deficit in hamstring strength (HSS) is related to an increased risk of suffering HSI [2–8]. Furthermore, deficits in HSS go along with a decrease in performance in running, sprinting and jumping tasks [9–13]. Therefore, the frequent assessment of HSS plays a crucial role in terms of athletes’ health and performance. Tests to assess HSS should not only be objective, valid and reliable; they should furthermore be easy and quick to perform to be implemented on a regular basis and should not put athletes’ health at additional risk. The assessment of HSS via single joint motorized gold standard dynamometry goes along with the downside of a high expenditure of time, cost and staff, and its immobility. For the sake of avoiding additional risk of HIS and hamstring muscle fatigue, isometric contractions are favored over eccentric contractions to assess HSS, as eccentric muscle actions lead to more extensive and longer lasting muscle damage than isometric contractions [14,15].

HSI most frequently occur during the late swing phase in sprinting, or the follow through after kicking a ball, where the knee is in a rather extended and the hip in a flexed position [16,17]. More than 80 % of HSI affect the m. biceps femoris [1,18] and the very same muscle is in its optimal force /torque:length ratio at positions of 15°-30° knee flexion [19,20]. Cohen et al. [21] also showed, that the hamstring muscle is able to generate its highest torque under 10° and 30° of knee flexion for eccentric and concentric contractions, respectively. They furthermore observed that eccentric hamstring muscle fatigue is the largest under 10° of knee flexion. These angular positions should be taken into consideration when designing tests to assess HSS with the goal of preventing HSI. Therefore, the objective and user-friendly assessment of HSS via mobile force plate or sphygomanometer under knee and hip flexion angles of 30° [20,22–26] or under 90° of hip and 20° of knee flexion [27], respectively, was proposed in the past. Assessing hamstring strength under a hip and knee flexion of 30° each [20,22–24] or 90° and 20° [27], respectively, led to a larger sensitivity to detect hamstring muscle fatigue than tests under 90° of hip and knee flexion, as proposed in an earlier study [28]. Also in terms of interday reliability, Cuthbert et al. [25] measured higher ICCs for assessing HSS under 30° than under 90° of knee and hip flexion, respectively. Read et al. [20] observed significantly higher biceps femoris activity (d = 1.19) under 30° of knee and hip flexion compared to 90° of knee and hip flexion, while no significant differences were present for the m. semitendinosus and m. gluteus maximus, respectively. On top of that, an increase in hip flexion goes along with an increase in muscle activity in the m. biceps femoris, m-semitendinosus, and m. gluteus maximus [29]. Hence, assessing HSS under 90° of hip flexion and 20° of knee flexion, as suggested in the 90:20 Isometric Posterior Chain Test (90:20 IPCT) [27], should be the favored position. Despite the promising results of the aforementioned studies, no validation of either of the tests against gold standard dynamometry has been carried out so far.

Read et al. [20], who assessed HSS under 30° of hip and knee flexion, observed that unprecise instruction led to different interpretations of how to perform the task, resulting in variations in horizontal force application during the execution of the test. While some of their participants applied anterior horizontal force onto the force plate, others tried to pull the force plate towards them, although subjects were instructed to only “push” vertically into the force plate. During piloting, we observed that during the 90:20 IPCT some participants did not only exert a vertical force onto the force plate, as instructed [27], but also applied anterior horizontal force at the same time, which may influence the peak vertical force component representing the main parameter. These differences in execution may have an influence on the vertical force as outcome parameter and account for interpersonal differences in HSS. We therefore added a new, more precise, instruction where the subjects were told to not only apply vertical but vertical *and* posterior horizontal force onto the force plate – i.e. pull the force plate towards them. We assumed that the original instruction of the 90:20 IPCT [27] rather targets hip extension capabilities of the posterior chain, whereas our new instruction is better suitable to represent knee flexion capacities and thus HSS.

Hence, this study aimed to compare the influence of two different verbal task instructions of the 90:20 IPCT on the interday reliability, validity and activation of the respective muscles. Our working hypothesis was that our new instruction shows equal or better reliability and leads to higher validity with gold standard dynamometry than the original instruction proposed by Matinlauri et al. [27]. Furthermore, we expected that a change of instruction leads to differences in muscle activation for the m. biceps femoris, m. semitendinosus, m. gluteus maximus, and m. rectus femoris during the 90:20 IPCT. These information can give important insights whether the 90:20 IPCT can be used as a cost and time efficient tool to assess HSS under the aspect of performance and HSI prevention, respectively.

## Methods

### Subjects

We recruited 23 (m: 12; w: 11) physically active participants (age: 27.9 ± 4.5 years; body mass: 70.8 ± 8.4 kg; height: 1.76 ± 0.06 m), who did neither have any physical impairments of the lower limbs nor back pain within the last three months. The subjects were informed about the purpose, methods and potential risks of the study and provided written consent in accordance with the Declaration of Helsinki. Ethics approval was obtained by the Technical University of Munich’s Ethics Committee (#2023-415-S-SB).

### Warm-up

Before the familiarization and the actual test days, all participants performed a standardized warm-up, which started with 5 mins of cycling on an indoor cycle ergometer (Puls Med 300, Dynavit, Kaiserslautern, GER) at 60-70 rpm and 100 W, followed by 2 mins at 150 W at the same cadence. After that, the subjects carried out three 30 m-sprints on an indoor sprint track at 60, 70 and 80 % of their subjective maximum sprint velocity, respectively, with 30 s break between each sprint. This was followed by two bunkie-tests (box height: 30 cm) of a duration of 20 s and 1 min break between the two tests.

### Experimental Setup

For the 90:20 IPCT, the unshod participants stood with their heels, buttocks, shoulders and head against the wall and kept their arms crossed at their chest, as this arm position provides the highest reliability [27]. Participants then placed one foot with the heel onto a three-axis force plate (FP4060-10-TM-2000, Bertec, Columbus, Ohio, USA), which was positioned on a height-adjustable table (Sympas STS JCHT35K28A, Assmann Büromöbel GmbH & Co., Melle, GER). With the help of a manual goniometer (Model 01135, Lafayette Instruments Co., Sagamore, Indiana, USA), the height of the table was electrically adjusted to the nearest centimeter so that the hip and knee flexion angles of the subjects could be set to 90° and 20°, respectively, with the ankle in a neutral position (Fig 1, left). The individual position of the heel on the force plate was marked with a piece of tape and noted down to ensure the same positioning throughout all trials and test days.

**Fig 1:**
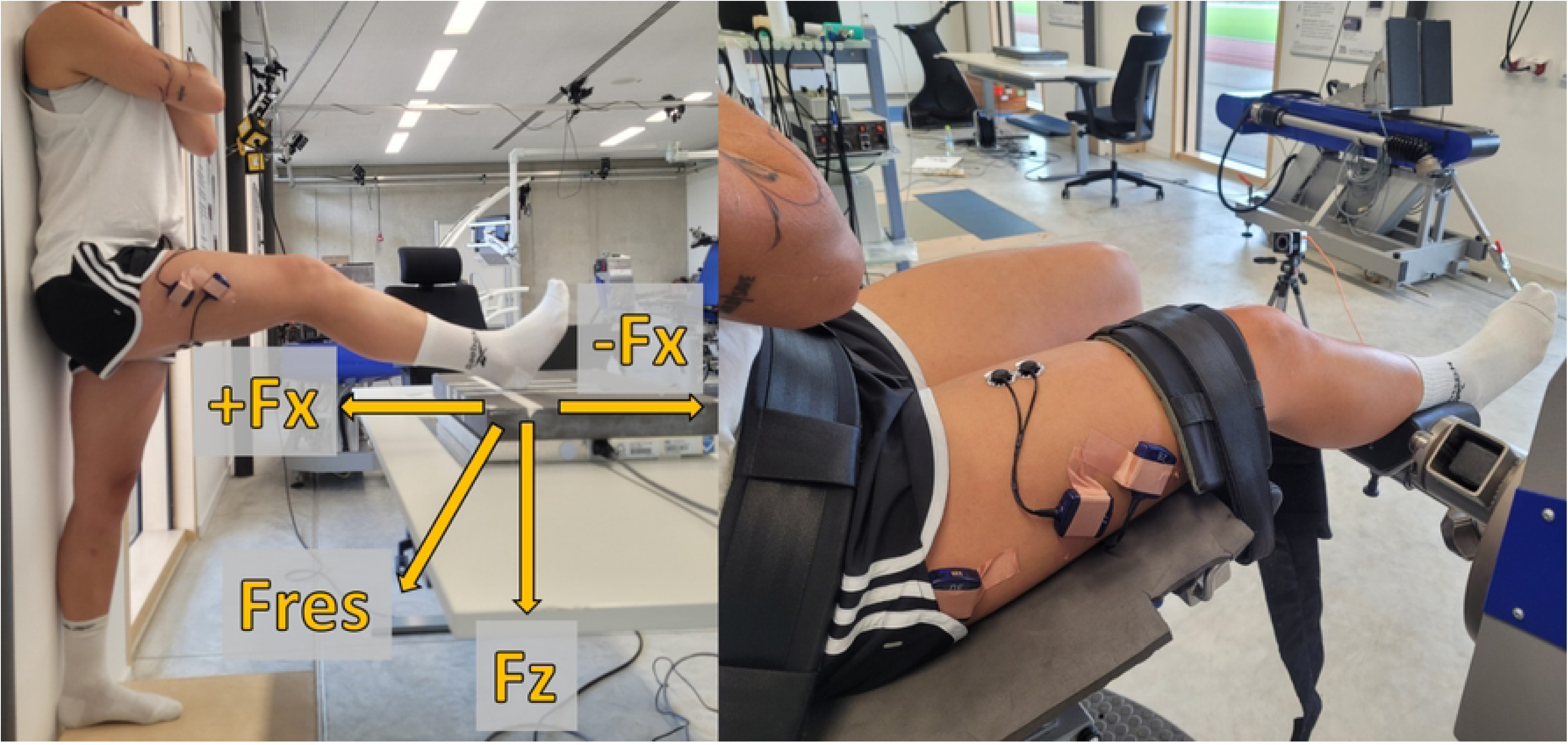
left: Set-up of the 90:20 IPCT; right: Set-up of isometric gold standard dynamometry. Fz: vertical force; +Fx: posterior horizontal force; -Fx: anterior horizontal force; Fres: resultant force.

The isometric gold standard measurements were carried out on a motor-driven dynamometer (IsoMed 2000, D.&R. Ferstl GmbH, Hemau, GER) in a sitting position, where the hip and knee angles were equivalent to those of the 90:20 IPCT. Therefore, the backrest and the seating surface were inclined to 70° and 20°, respectively. The axis of rotation of the dynamometer was aligned with the axis of rotation of the knee joint and the dynamometer arm was positioned with the help of a manual goniometer, so that the knee was flexed to 20°. For the measurements, the subjects were fixed to the device with a waist belt, shoulder pads, and a strap around the lower thigh to avoid unwanted movement during the contraction (Fig 1, right). To account for a possible shift of the knee axis of rotation during isometric contraction (e.g., soft tissue compression), the alignment of the axes was done while the participant was contracting.

During the 90:20 IPCT, we measured muscle activities of the m. rectus femoris, m. biceps femoris (long head), m. semitendinosus, and m. gluteus maximus via surface EMG (Myon 320, Myon AG, Schwarzenberg, CH). Therefore, the skin was shaved, roughened and cleaned in the first step. Second, electrodes (H124SG, Cardinal Health, Dublin, Ohio, USA) were attached to the subjects according to the SENIAM guidelines [30].

All devices were connected to a 16-bit A/D converter (NI USB-6218, National Instruments, Austin, Texas, USA), and data were recorded with ProEMG 2.1.0.1 (Prophysics AG, Kloten, CH) at a sampling frequency of 1000 Hz.

### Experimental Protocol

Participants took part in the measurements on two separate test days (T1 and T2) with two to four days in between. The time of day was kept identical for the test days to control for circadian rhythm. We also asked the subjects to refrain from strenuous physical activity 48 h hours prior to the measurements; furthermore, we encouraged them to pursue the same daily routine on and before each test day. Adherence to the specifications was verbally queried on the day of the test.

Two to five days before T1, the subjects received a familiarization session. Familiarization was successful when the participants carried out three consecutive trials within a range of ± 5 % of the maximum force/ torque at the respective tests. In addition, for the vertical-horizontal instruction of the 90:20 IPCT, the subjects had to pull the force plate towards them with a posterior horizontal force of at least 10 N. The familiarization session was also used to set-up the devices according to the subjects’ anthropometry to ensure the same positioning throughout the two sessions. All measurements were carried out with the dominant leg, which was defined as the leg the subjects would use for kicking a ball (left-dominant: n = 8; right-dominant: n = 15).

After the warm-up on the respective test day, we randomized the order of the tests, where randomization was done separately for T1 and T2. For the 90:20 IPCT, participants received the original instruction as proposed by Matinlauri et al. [27] to “*Exert maximal force vertically into the plate”* (vertical instruction: V) and our new instruction “*Exert maximal force vertically into the plate and pull the plate towards you”* (vertical-horizontal instruction: VH). At the beginning of the trials of the 90:20 IPCT, subjects were asked to relax in the respective position for 2 s to acquire the horizontal gravitational forces of the leg. Data acquisition with the dynamometer was carried out under the instruction *“Flex your knee joint as hard as you can.”* During all test trials, participants exerted maximal force/ torque for about 5 s – or as long as the force/ torque curve inclined – and were maximally verbally encouraged. Data acquisition on the two test days was carried out by the same examiner to avoid interpersonal effects on the reliability. We also asked the subjects at T1 to redraw the positions of the EMG electrodes with permament marker to ensure their identical positioning at T2. After each trial, the test was alternated preceding a 60 s break. The subjects carried out a total of three trials per test on each of the test days.

### Data Analysis

Using a custom-written Matlab script (Version R2020b, The MathWorks, Inc., Natick, Massachusetts, USA), EMG data were centered, and filtered with a 4^th^ order zero-lag Butterworth bandpass filter with cut-off frequencies of 10 and 499 Hz, respectively. After rectification of the EMG data, both EMG and force/ torque data were smoothed with a 250 ms moving average to avoid latency between the kinetic and the EMG data. The horizontal gravitational forces of the leg (1 s-mean under relaxed state) at the 90:20 IPCT were subtracted from the exerted horizontal force to define whether the subjects applied anterior or posterior horizontal forces onto the force plate, respectively. Parameters of interest for the 90:20 IPCT were peak vertical force Fz under both, the original vertical [27] and our new vertical-horizontal instruction (Fz_V and Fz_VH). We furthermore extracted the anterior-posterior horizontal force Fx at the moment of the peak Fz for either instruction (Fx@Fz_V and Fx@Fz_VH) as an additional parameter. For the vertical-horizontal instruction (VH) only, the peak resultant force (Fres_VH) was calculated when the plate was pulled towards the subjects (Fx > 10 N). We further multiplied the peak force component that pointed orthogonally to the shank (F_ortho_VH) with the length of the subjects’ shank (distance from lateral epicondyle to lateral malleolus plus mean distance from lateral malleolus to the floor) [m] to transform this force into torque (M_F_ortho_VH) [Nm].

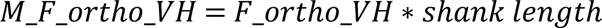

F_ortho_VH was calculated via the following formula (Fig 2 for angles and force orientations):

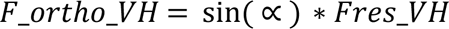

**Fig 2:**
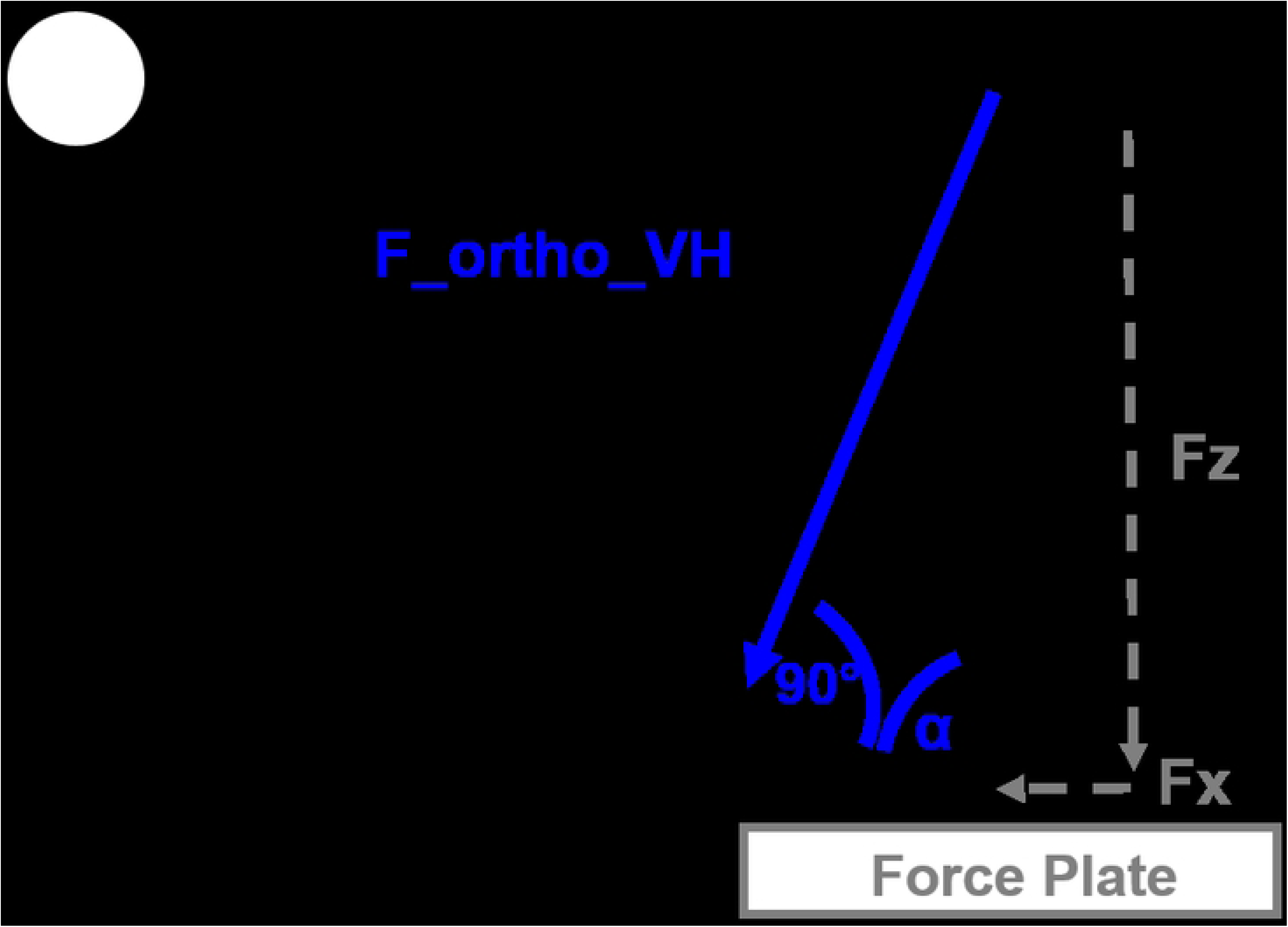
Definition of angles and force vectors for the calculation of the force vector orthogonally to the subject’s shank. Fz: vertical force component of resultant force under vertical-horizontal instruction; Fx: posterior horizontal force component of resultant force under vertical-horizontal instruction; Fres_VH: resultant force under vertical-horizontal instruction; F_ortho_VH: force component of resultant force under vertical-horizontal instruction orthogonally to the shank.

, where Fres_VH and its corresponting angle β in relation to the force plate were calculated as:

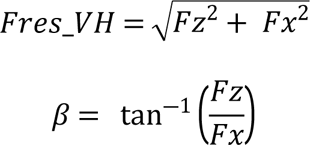

To be able to finally calculate F_ortho_VH via the sine of α, we calculated this angle via:

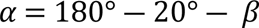

For the isometric dynamometry measurements, we extracted the peak torque of the three trials (M). An overview of the test order and the parameters derived from the respective tests is given in Fig 3.

**Fig 3:**
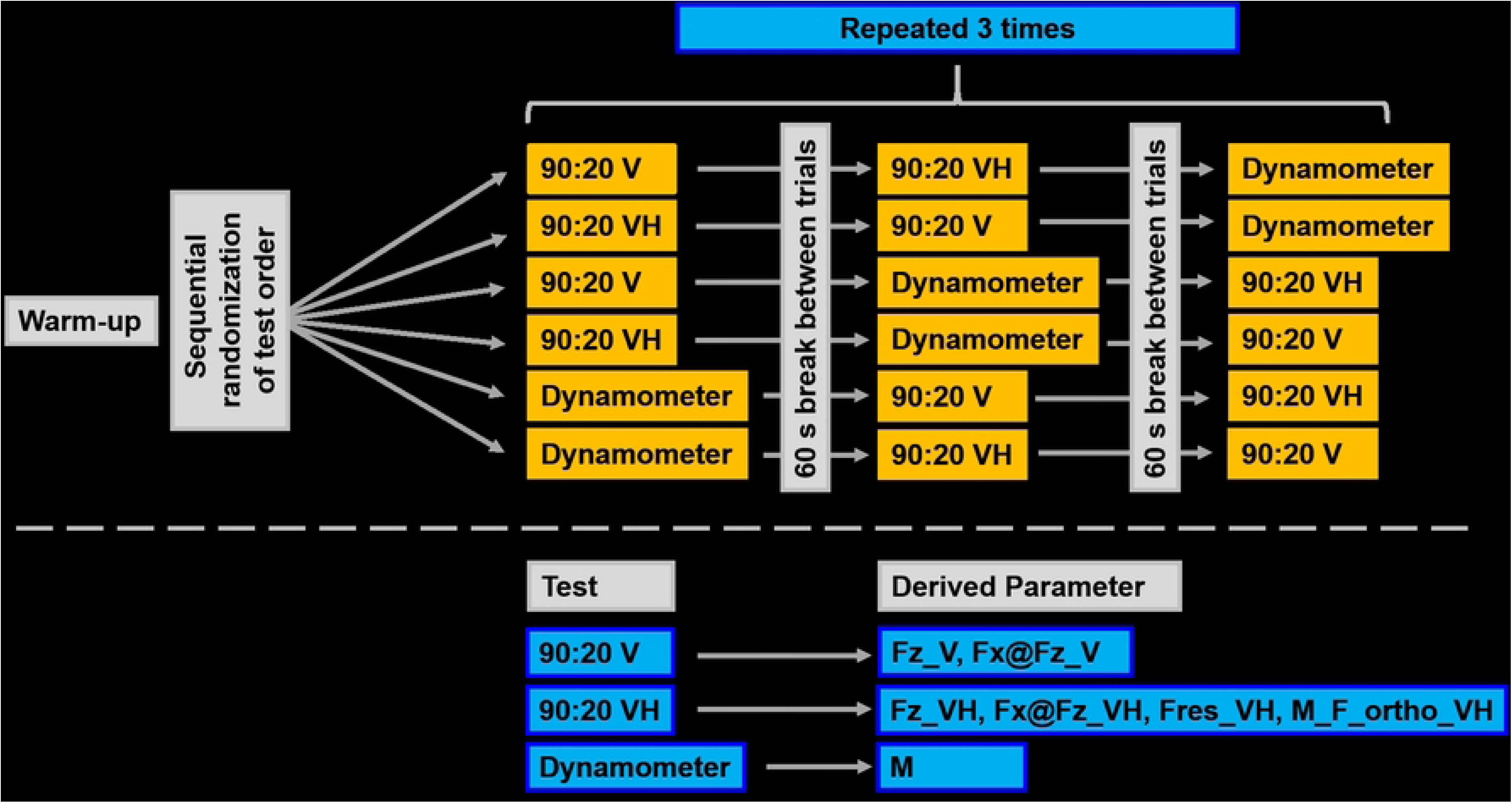
Overview of test order and measured parameters. 90:20 V: 90:20 Isometric Posterior Chain Test under vertical instruction; 90:20 VH: 90:20 Isometric Posterior Chain Test under vertical-horizontal instruction; Dynamometer: isometric gold standard dynamometry; Fz_V: peak vertical force under vertical instruction; Fx@Fz_V: horizontal force at the moment of peak vertical force under vertical instruction; Fz_VH: peak vertical force under vertical-horizontal instruction; Fx@Fz_VH: horizontal force at the moment of peak vertical force under vertical-horizontal instruction; Fres_VH: peak resultant force under vertical-horizontal instruction; M_F_ortho_VH: torque calclulated from force vector of resultant force orthogonally to the shank under vertical-horizontal instruction multiplied with shank length; M: peak torque.

For a further understanding of the forces and the corresponding extracted parameters, an exemplary force-time graph under vertical-horizontal instruction is provided in Fig 4.

**Fig 4:**
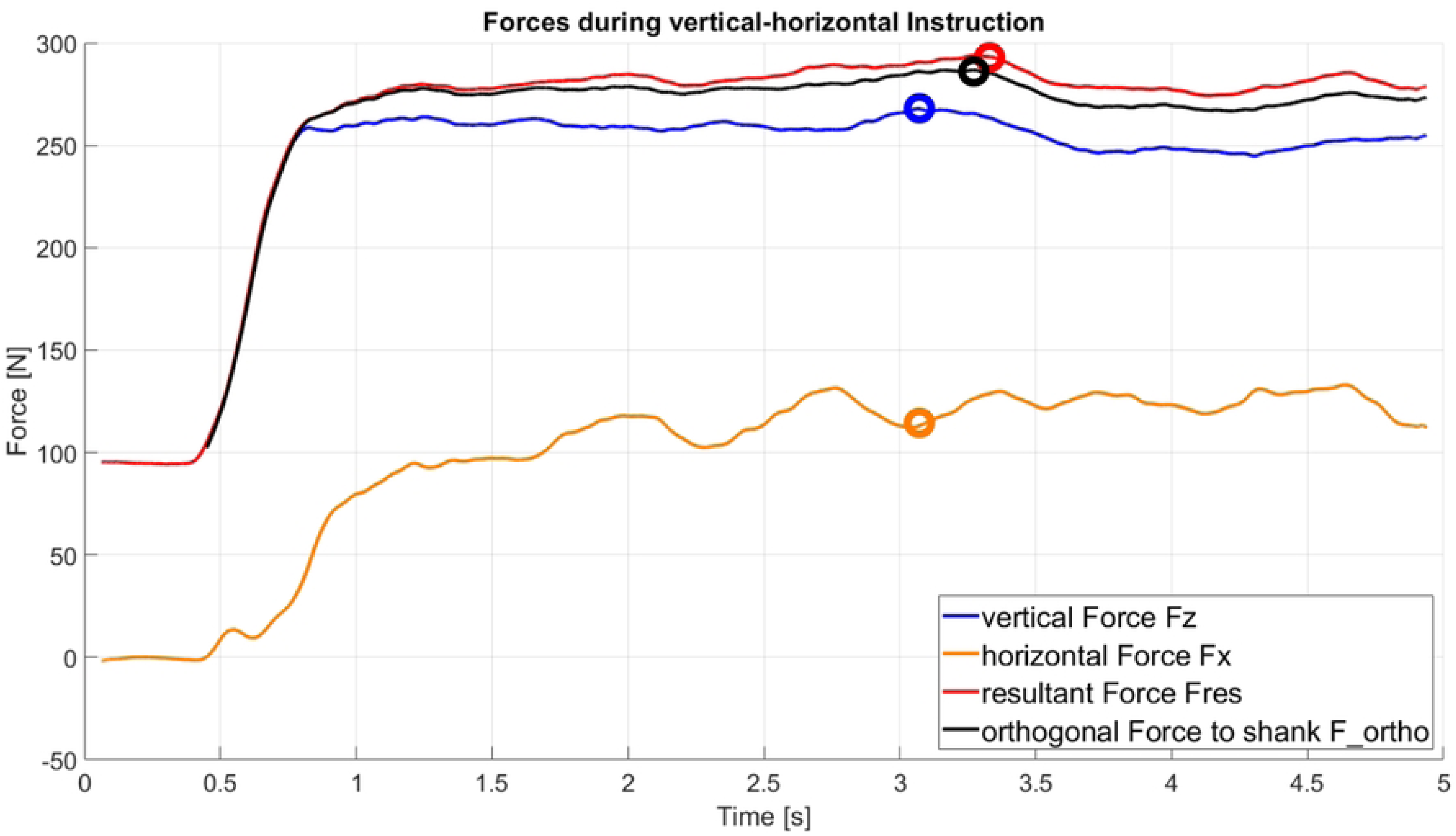
Exemplary force-time curve under vertical-horizontal instruction. Circles depict the maximum value of the respective parameter or, for Fx, the corresponding value at the moment of peak Fz.

To assess the activity of the aforementioned muscles, we calculated the mean EMG of the period when the force/ torque was above 95 % of the peak value during the respective trial. The corresponding EMG data of the trial with the largest kinetic value were included for statistical analysis of the muscle activities. The main purpose of this study was to evaluate whether the vertical-horizontal instruction (VH) and its parameters lead to a difference in muscle activities compared to the original vertical instruction (V). Therefore, the muscle activities of the m. biceps femoris, m. semitendinosus, m. gluteus maximus and m. rectus femoris at peak vertical force under the original instruction (Fz_V) served as reference (= MVC).

### Statistical Analysis

To assess the inter-day reliability of the described parameters, we calculated intraclass correlation coefficients (ICC3,1). Interpretation of ICCs took place according to Koo and Li [31] where ICC values and their corresponding 95 % confidence intervals (95%CI) of < 0.5, 0.5-0.75, 0.75-0.9, and > 0.9 are regarded as poor, moderate, good and excellent, respectively. As comparable literature on the assessment of HSS does not only use ICCs but also coefficients of variation (CV) as a measure of reliability, we also calculated the respective CVs for a better interpretation of our data. Acceptable reliability was given for CVs < 15 % [32].

We secondly checked for the validity of the different instructions of the 90:20 IPCT by calculating correlation coefficients (Pearson’s r or Spearman’s rho in the case of violation of normality (Shapiro-Wilk test)) between the parameters of the 90:20 IPCT with isometric gold standard dynamometry for each test day. Correlation coefficients of less than 0.2 were classified as very weak, whereas a weak, moderate, strong and very strong agreement was given for values from 0.2-0.39, 0.4-0.59, 0.6-0.79, and > 0.8, respectivley [33].

As Read et al. [20] observed a variance in horizontal force due to unprecise instruction, we furthermore calculated paired samples t-tests to assess whether the peak vertical force changes with a different instruction (Fz_V vs. Fz_VH). We did the same with the horizontal force at the moment of peak vertical force (Fx@Fz_V vs. Fx@Fz_VH) to see whether the horizontal forces statistically differ from each other with a change of instruction. On top of that, calculated correlation coefficients between peak vertical force (Fz) and the corresponding horizontal force at the moment of peak vertical force (Fx@Fz) for both instructions to see, if there is a relationship between these two force components.

Descriptives of the EMG activities (% of deviation from reference) are given in mean and standard deviation, or median and interquartil range (IQR), where applicable. Eventually, EMG activities of the examined muscles were compared to the muscle activities of the peak vertical force of the 90:20 IPCT under the original vertical instruction (Fz_V). We did so by conducting two tailed one sample t-tests – or Wilcoxon signed-rank tests in the case of violation of normality – against the reference values (Fz_V = 100 %) and calculated the according effect sizes (Cohen’s d or matched rank biserial correlation (rbc), respectively). Effect sizes are regarded as small, medium or large for Cohen’s d values > 0.20, > 0.50, and > 0.70, respectively [34]. Level of significance was set at α = 0.05 for all tests; for the one sample t-tests of the EMG activities, however, we made a Bonferoni adjustement of our alpha level according to the number of tests. All statistical calculations were carried out with JASP 0.16.4 [35].

## Results

### Reliability

In terms of inter-day reliability, isometric gold standard dynamometry delivered excellent values (ICC: 0.96; 95%CI: 0.90–0.98). M_F_ortho_VH, Fz_V and Fres_VH showed comparably good to excellent reliability with ICCs of 0.94 (95%CI: 0.87-0.98), 0.91 (95%CI: 0.80-0.96) and 0.89 (95%CI: 0.75-0.95), respectively, whereas reliability for Fz_VH could be regarded as moderate to excellent (ICC: 0.87; 95%CI: 0.72-0.94).

The calculation of the corresponding coefficients of variation (CV) of the aforemetioned parameters resulted in comparibly acceptable reliability for M (CV: 4.7 %), M_F_ortho_VH (CV: 4.7 %), Fz_V (CV: 5.6 %), Fres_VH (CV: 4.1 %), and Fz_VH (CV: 4.7 %), respectively.

### Validity

In terms of agreement of the parameters of the 90:20 IPCT with isometric gold standard dynamometry, M_F_ortho_VH showed a very strong correlation with r = 0.83 and r = 0.87 (p < 0.001, each) for T1 and T2, respectively. Fres_VH also correlated very strongly (T1: r = 0.81, p < 0.001) and strongly (T2: r = 0.80, p < 0.001) with gold standard. Both Fz_VH (T1: r = 0.80, p < 0.001; T2: r = 0.73; p < 0.001) and Fz_V (T1: r = 0.69, p < 0.001; T2: r = 0.70, p < 0.001) showed strong agreement with the peak torque of gold standard dynamometry (Fig 5).

**Fig 5:**
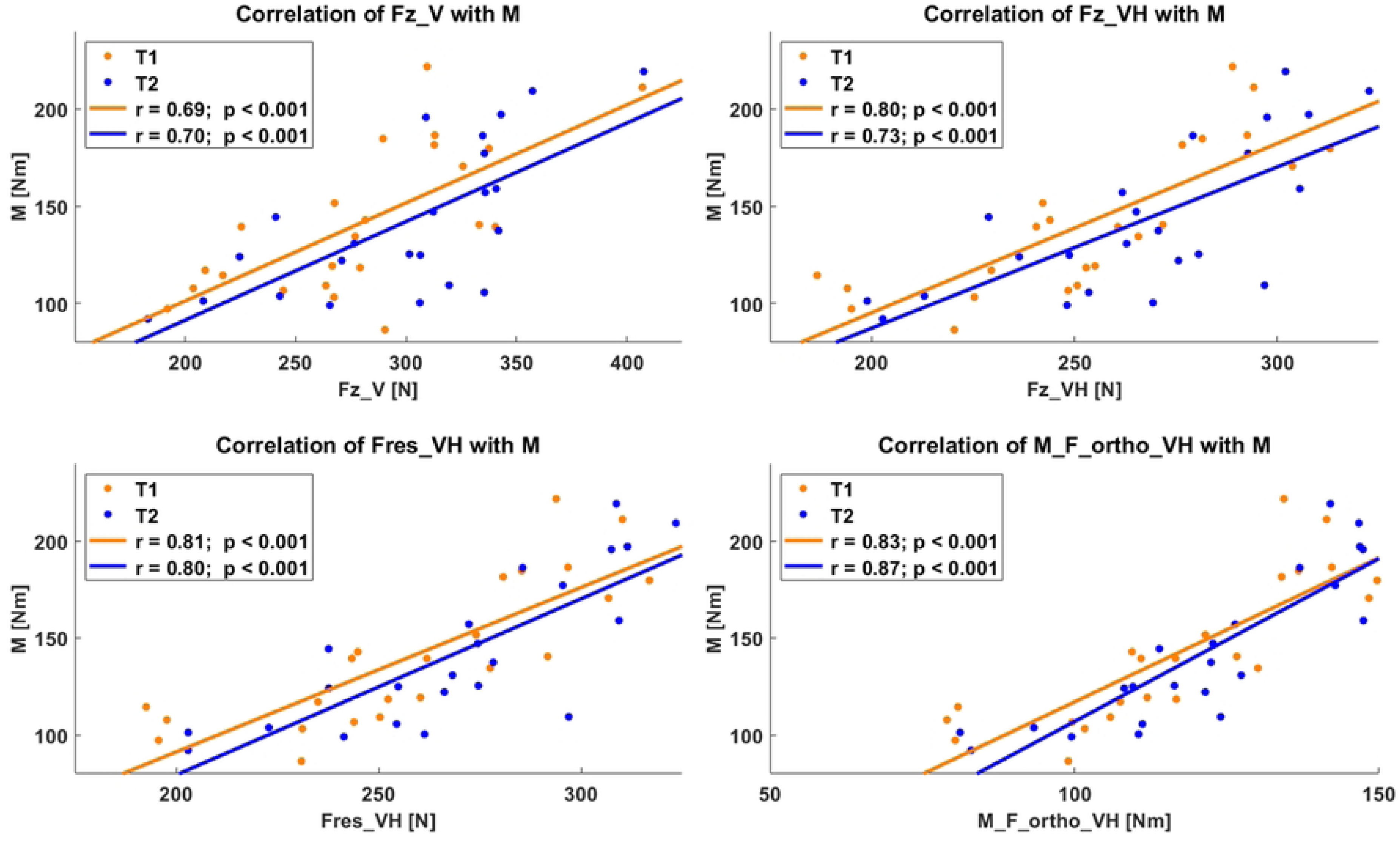
Agreement of the various parameters with gold standard dynamometry (M). Fz_V: peak vertical force under vertical instruction; Fz_VH: peak vertical force under vertical-horizontal instruction; Fres_VH: peak resultant force under vertical-horizontal instruction; M_F_ortho_VH: moment calculated via shank length multiplied by force component orthogonally to the shank.

### Influence of Instruction on identical Parameters

Comparing vertical force under either instruction (Fz_V and Fz_VH) shows that vertical force under vertical instruction is significantly higher than the very same parameter under vertical-horizontal instruction for either measurement days with a strong effect size, each (T1: Fz_V: 281 ± 52 N vs. Fz_VH: 254 ± 35 N, p < 0.001, d = 0.87; T2: Fz_V: 300 ± 54 N vs. Fz_VH: 266 ± 34 N, p < 0.001, d = 1.10). For the corresponding horizontal forces at peak vertical force under vertical instruction (Fx@Fz_V), -38 ± 30 N and -45 ± 39 N were measured for T1 and T2, respectively. Under vertical-horizontal instruction, a horizontal force at peak vertical force (Fx@Fz_VH) of 32 ± 23 N was observed for T1, whereas T2 showed a median horizontal force of 38 IQR 50 N). The differences between the two instructions for horizontal force at peak vertical force were significant for both T1 (p < 0.001, d = -1.78) and T2 (p < 0.001, d = -1.50), respectively (see Fig 6).

**Fig 6:**
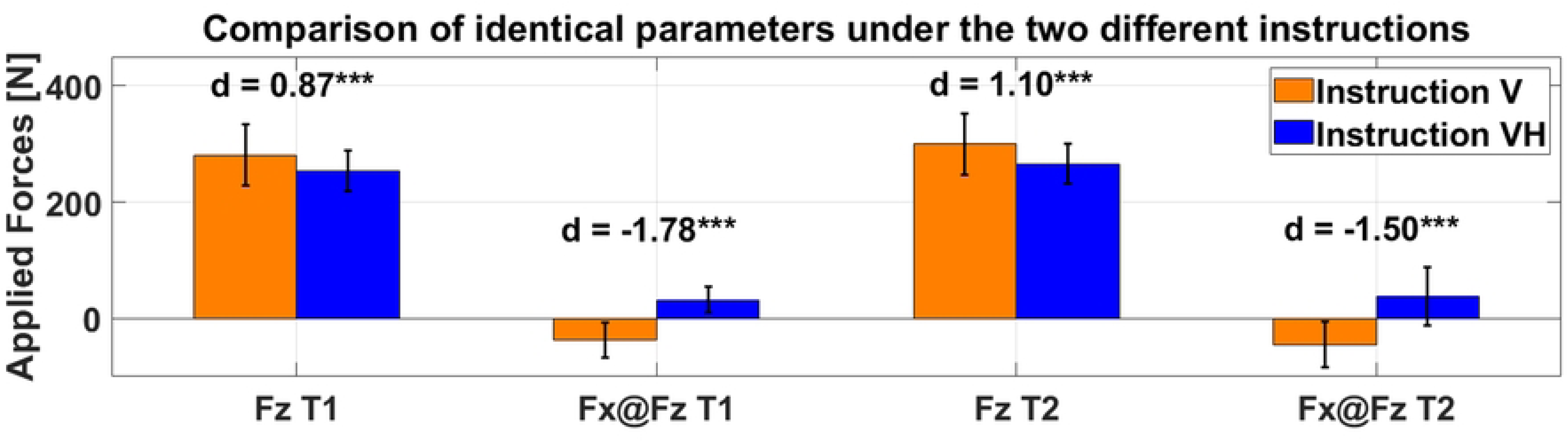
Comparison of vertical instruction (orange) with vertical-horizontal instruction (blue) for the parameters peak vertical force (Fz) and horizontal force at the moment of peak vertical force (Fx@Fz). ***: p < 0.001; **: p < 0.01; *: p < 0.05.

Furthermore, there was a moderate to strong negative correlation between peak vertical force and the corresponding horizontal force at the moment of peak vertical force under vertical instruction (Fz_V and Fx@Fz_V) (T1: r = -0.54; p = 0.008; T2: r = -0.66; p < 0.001), whereas no relationship between these two parameters was present under vertical-horizontal instruction (Fz_VH and Fx@Fz_VH) (T1: r = 0.15; p = 0.50; T2: r = 0.13; p = 0.57).

### Muscle Activity

For the analysis of muscle activities between the two instructions and the corresponding parameters, we considered the original reference parameter Fz_V, the alternative parameter that can be used with a single axis force plate (i.e. Fz_VH), and the parameter derived from vertical *and* horizontal forces that shows the best agreement with gold standard dynamometry (i.e. M_F_ortho_VH). We excluded the m. rectus femoris from statistical analysis due to an insufficient signal to noise ratio of < 3 [36] during all of the tests. This indicates, that the m. rectus femoris did not actively contribute to either of the tests. For the further statistical evaluation of the remaining muscle activities, we had to exclude one subject for each of the muscles and comparisons due to interferred EMG signals. For the m. biceps femoris, we observed a slight increase of muscle activity during peak vertical force under vertical-horizontal instruction (Fz_VH) compared to bare vertical instruction (Fz_V) of the 90:20 IPCT. These differences, however, were only marginal (T1: +5.7 ± 19.4%, p = 0.184; T2: +4.3 ± 17.8 %, p = 0.273) and not statistically significant. The same applies for the muscle activity of the m. biceps femoris and the moment of the force component orthogonally to the shank under vertical-horizontal instruction (M_F_ortho_VH) (T1: +5.5 ± 18.0 %, p = 0.171; T2: +6.4 ± 16.6 %, p = 0.084, no tendency due to Bonferroni α-adjustment). No statistically significant differences were observed for the muscle activation of the m. semitendinosus between the different parameters of the 90:20 IPCT (Fz_VH: T1: +9.0 ± 28.3 %, p = 0.150; T2: -1.8 ± 12.1 %, p = 0.493; M_F_ortho_VH: T1: +8.5 ± 27.8 %, p = 0.165; T2: +2.6 ± 12.9 %, p = 0.348). Muscle activities of the m. gluteus maximus under vertical-horizontal instruction did not show significant differences compared to the muscle activity of the original parameter Fz_V (Fz_VH: T1: -17.2 ± 48.6 %, p = 0.503; T2: -3.0 ± 27.6 %, p = 0.601; M_F_ortho_VH: T1: - 15.9 IQR 41.2 %, p = 0.483; T2: -3.2 IQR 26.8 %, p = 0.571) (see Fig 7).

**Fig 7:**
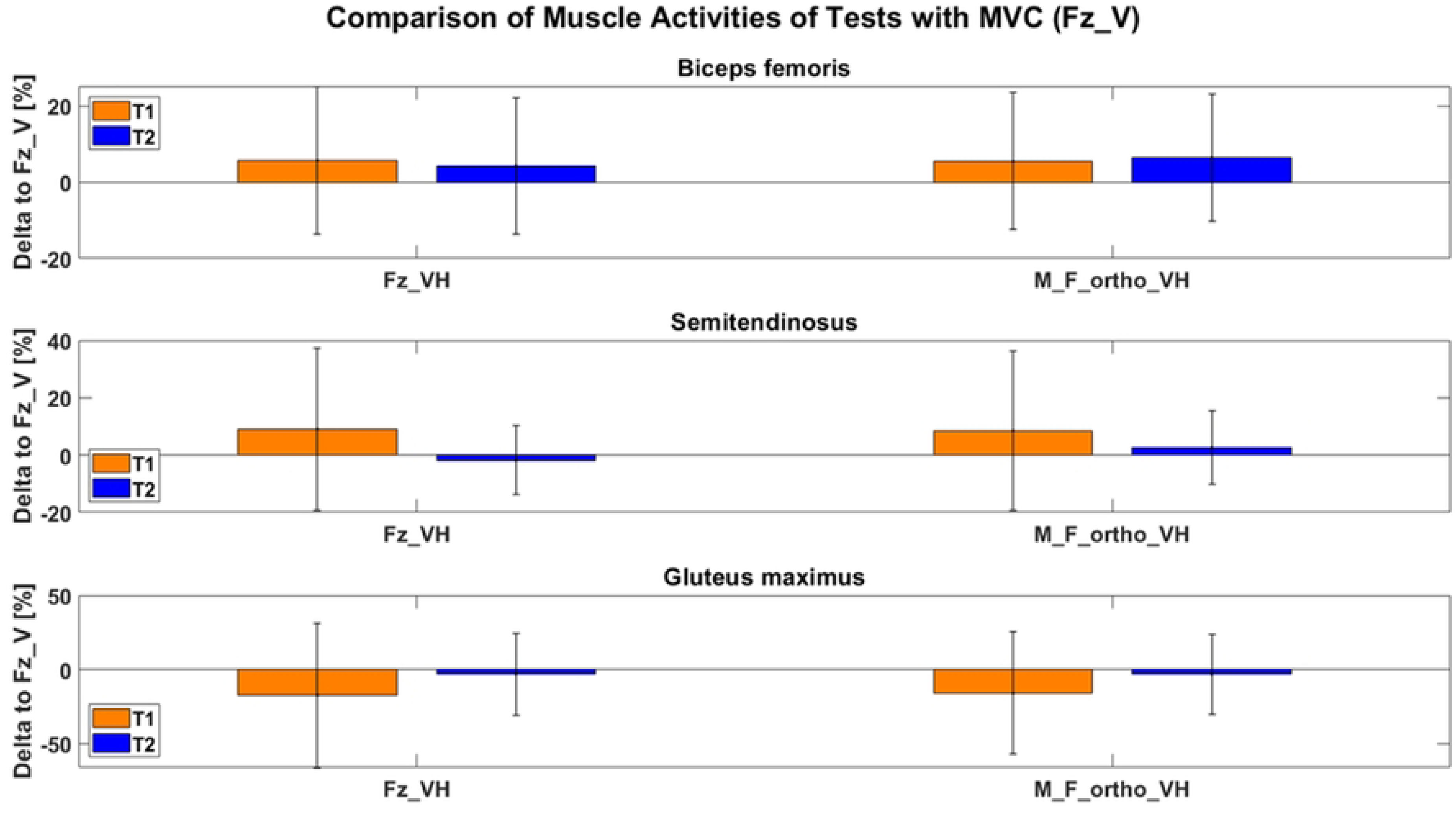
Comparison of muscle activities of the different tests with the muscle activities of vertical force under vertical instruction (= reference). Fz_VH: peak vertical force under vertical-horizontal instruction; M_F_ortho_VH: moment calculated by shank length multiplied by peak force component orthogonally to the shank.

## Discussion

This paper compared the original outcome parameter Fz_V of the 90:20 IPCT with parameters of this very test under a different instruction. Reliability, agreement with isometric gold standard dynamometry, and the differences in activities of the involved muscles were objects of interest. We furthermore tested if a change of instruction (V vs. VH) has an influence on the magnitude of the outcome parameter of bare vertical force (Fz_V) and its corresponding horizontal force (Fx@Fz).

Regarding reliability, the force parameters of the 90:20 IPCT, acquired in this study (Fz_V, Fz_VH and Fres_VH), showed similar reliability. Calculation of the applied torque (M_F_ortho_VH) during the 90:20 IPCT via multiplying the maximum force component orthogonally to the shank (F_ortho_VH) and the shank length delivers even better reliability than the force parameters of the 90:20 IPCT with an ICC of 0.94 and comes closest to the values of isometric single joint gold standard dynamometry reaching an ICC of 0.96. McCall et al. [23] observed moderate to good reliability for the dominant leg (ICC: 0.86; 90%CI: 0.69-0.94) and good to excellent reliability for the non-dominant leg (ICC: 0.93; 90%CI: 0.84-0.97) concerning peak vertical force under 30° of hip and knee flexion. Similarly good ICCs were found in Nedelec et al. (ICCs: 0.80-0.88), who also assessed HSS under the same knee and hip angles. Cuthberg et al. [25], who also carried out the same test, only measured poor to good reliability for the left leg (ICC: 0.76; 95%CI: 0.29-0.82), whereas moderate to excellent reliability was observed for the right leg (ICC: 0.86; 95%CI: 0.64-0.94). Using the coefficient of variation as a measure of reliability, all of the parameters of the 90:20 IPCT in this study generate similar values (CVs: 4.1 %-5.7 %) and can thus be considered equally suitable to assess HSS reliably [32]. Matinlauri et al. [27] reported low relative inter-day variability (CV < 8.1 %) for the vertical force under vertical instruction (Fz_V) of the 90:20 IPCT. For the tests under 30° of knee and hip flexion, Constantine et al. [24] observed a similarly low variablity with a mean coefficient of variation of 7.5 %. The parameters assessed in our study are therefore in line with comparable literature and show equally good or even better reliability.

Regarding validity, the original instruction and parameter of the 90:20 IPCT (Fz_V), assessed in this study shows a strong agreement with gold standard isometric dynamometry for both T1 (r = 0.69) and T2 (r = 0.70). The parameters under vertical-horizontal instruction, however, all show better agreement with gold standard than the original parameter. Vertical force under vertical horizontal instruction (Fz_VH) delivers very strong to strong agreement with gold standard (T1: r = 0.80 and T2: r = 0.73) whereas the resultant force under vertical-horizontal instruction (Fres_VH) leads to a further, very strong agreement with isometric dynamometry for both measurement days (r = 0.81 and r = 0.80). The applied torque (M_F_ortho_VH) during the 90:20 IPCT shows the best agreement with gold standard among the examined parameters with r = 0.83 and r = 0.87 for T1 and T2, respectively. Squaring the Pearson’s r coefficients of the original parameter (Fz_V) and the parameter of the vertical-horizontal instruction, which shows the best agreement with gold standard (i.e. M_F_ortho_VH), results in a mean r^2^ of 0.48 for Fz_V and r^2^ of 0.72 for M_F_ortho_VH. Hence, compared to Fz_V, M_F_ortho_VH explains 50 % more variance, or 24 percentage points, respectively, and therefore better represents HSS assessed with the 90:20 IPCT than the original instruction and parameter. There is no study comparing the outcome parameters of the 90:20 IPCT with gold standard dynamometry, yet. Nedelec et al. [22] show a moderate correlation of HSS assessed under 30° of knee and hip flexion of r = 0.53 (95%CI: -0.03-0.84; p = 0.06) with gold standard dynamometry, whereas HSS under hip and knee flexion angles of 90° delivered a very strong agreement (r = 0.81; 95%CI: 0.48-0.94; p = 0.001). In their study, however, they only used the vertical instruction for either angular conditions. Vertical-horizontal instruction and M_F_ortho_VH, as assessed in this study, therefore generate increased agreement to gold standard compared to tests only considering the vertical force component under vertical instruction (Fz_VH) under 30° or 90° of hip and knee flexion [22] and 90° and 20°, respectively.

The differences, we measured in peak vertical force between the two instructions (Fz_V and Fz_VH) show that – as assumed by Read et al. [20] – the instruction or rather the interpretation of the instruction by the respective participant has a strong influence on the outcome parameter Fz. For peak vertical force, the vertical instruction (Fz_V) leads to significantly higher forces than the identical parameter under vertical-horizontal instruction (Fz_VH). Consequently, differences between athletes’ HSS measured via bare vertical force under vertical instruction (Fz_V) of the 90:20 IPCT are not necessarily attributable to differences in HSS, but can be caused by a different execution of the rather unprecise vertical instruction, as it leaves more room for individual interpretation. Alike the vertical force, there were also significant differences between the horizontal force at the moment of peak vertical force, where under vertical instruction the force plate was significantly more pushed away (- Fx) from the subjects than under vertical-horizontal instruction, where the force plate was pulled (+Fx) towards the subjects’ body, which emphasizes the correct execution of the vertical-horizontal instruction and its practicability. On top of that, we observed a moderate to strong significant correlation between between the peak vertical force under vertical instruction (Fz_V) and the corresponding horizontal force component (Fx@Fz_V), which indicates that these force components are not independent of each other. This observation backs our and Read et al.’s [20] initial assumption that an instruction, leading to anterior horizontal force, rather adresses a “hip extension dominant production of force”, as the heel on the force plate acts as a counter bearing that experiences an anterior shear force when extending the hip. Consequently, the more hip extension is carried out, the more vertical force is applied and the more anterior horizontal shear force is generated, eventually. This might explain the fact that Nedelec et al. [22] measured a worse validity for 30° of knee and hip flexion than for 90° of knee and hip flexion, as the rather extended hip and knee leaves more room for individual interpretation and the application of more anterior force due to the rather unprecise vertical instruction, whereas single joint gold standard measurements rather required knee flexion capacities. This can also explain why in our study Fz_V showed less agreement with isometric single joint gold standard dynamometry than the parameters calculated under vertical-horizontal instruction. As opposed to the vertical instruction, no correlation was present between peak vertical force (Fz_VH) and horizontal force at the moment of peak vertical force (Fx@Fz_VH) under vertical-horizontal instruction, which means that neither of these force component is affected by the other.

Changes in muscle activitiy of the different parameters of the 90:20 IPCT under vertical-horizontal instruction were not significantly different from the reference (Fz_V) for neither of the muscles or test days. Therefore, our hypothesis that the m. biceps femoris, m. semitendinosus, and m. gluteus maximus show a different activation under vertical-horizontal instruction of the 90:20 IPCT compared to the original parameter (Fz_V) has to be rejected. The low maximum signal to noise ratio (< 3) of the m. rectus femoris shows that this muscle does not actively contribute to either of the tests. Furthermore, this finding indicates that the anterior horizontal force observed under vertical instruction (Fx@Fz_V) is not actively caused by the m. rectus femoris, but the results of the heel acting as a counter bearing during hip extension.

## Conclusion

Based on the results of this study, we recommend to *only* use the vertical-horizontal instruction and the calculation of the torque generated onto the force plate (M_F_ortho_VH) when assessing HSS with the 90:20 IPCT, as this parameter and instruction can explain 50 % more variance (r^2^ = 0.72) compared to the original parameter and instruction (Fz_V) with an r^2^ = 0.48. Furthermore, this parameter shows the highest reliability of the examined parameters of the 90:20 IPCT. To be able to acquire this parameter, however, a multi-axis force plate is required; besides this, this parameter and instruction do not require further additional equipment or time, except the once-only measurement of the shank length. Anyhow, where only a single-axis force plate is available, vertical force under vertical-horizontal instruction (Fz_VH) also delivers promising results and shows better agreement with isometric dynamometry than vertical force under vertical instruction (Fz_V); this, however happens at the cost to a slightly worse reliability.

To sum up, the 90:20 IPCT under our new vertical-horizontal instruction and the torque generated onto the force plate (M_F_ortho_VH) as outcome parameter is a reliable, valid and objective tool to assess isometric HSS. Its time efficiancy further emphasizes its potential assessing HSS on a regular basis to evaluate performance or return to sport readiness, or prevent HSI.

Future research on the optimized 90:20 IPCT should be made to find out, how this very test can be used to predict, or prevent HSI, respectively. Here, a cut-off value of relative torque that indicates an increased risk of HSI would be desirable for practitioneers and athletes to make use of this test to a full extent.

## Supporting Information

**S1 File. Raw Data.** (XLSX)

**S2 File. Table of processed Parameters for statistical Analysis.** (XLSX)

